# Targeting ALC1 can safely expand the therapeutic utility of PARP inhibitors across high-grade serous ovarian cancers

**DOI:** 10.64898/2025.12.04.692458

**Authors:** Lindsey N. Aubuchon, Deanna H. Wong, Natasha M. Ramakrishnan, Samantha F. Greenberg, Rohan Reddy, Elena Lomonosova, Amanda Compadre, Kathleen E. Jackson, Danielle Kemper, Vrutti Mehta, Kellen Zoberi, Dineo Khabele, Elizabeth L. Christie, Mary M. Mullen, Priyanka Verma

**Affiliations:** Division of Oncology, Department of Medicine, Siteman Cancer Center, Washington University School of Medicine; St. Louis, MO, 63110, USA; Cancer Biology Graduate Program, Washington University School of Medicine; St. Louis, MO, 63110, USA; Division of Gynecologic Oncology, Department of Obstetrics and Gynecology, Siteman Cancer Center, Washington University; St. Louis, Missouri, USA; Center for Reproductive Health Sciences, Department of Obstetrics and Gynecology, Siteman Cancer Center, Washington University School of Medicine; 4911 Barnes Jewish Hospital Plaza, St. Louis, Missouri 63110, USA; Sir Peter MacCallum Department of Oncology, The University of Melbourne; Parkville, Victoria, Australia

## Abstract

Poly (ADP-ribose) polymerase inhibitors (PARPi) are approved for homologous recombination-deficient (HRD) high-grade serous ovarian cancers (HGSOCs), but their long-term effectiveness is limited by the emergence of resistance and hematological toxicity. Moreover, PARPi are largely ineffective in HR-proficient HGSOCs, particularly tumors with *CCNE1* amplification, which exhibit marked therapeutic resistance and currently lack effective treatment options. Loss of a chromatin remodeling enzyme, Amplified in Liver Cancer 1 (ALC1), has been shown to enhance PARPi sensitivity. However, the clinical contexts in which ALC1 targeting will be clinically meaningful remain elusive. Here we demonstrate that ALC1 loss enhances PARPi sensitivity across HRD and *CCNE1*-amplified serous ovarian cancer lines, xenograft and patient-derived cells. ALC1 depletion can overcome clinically relevant mechanisms of PARPi resistance while having minimal effects in *BRCA*-wild-type or heterozygous non-cancerous cells. Consistent with this therapeutic safety, PARPi sensitivity upon ALC1 loss can be reliably predicted by the endogenous levels of phospho-T21 RPA2, a marker for replication stress which is typically higher in ovarian cancer cells compared to their normal counterparts. Together, our studies define the clinical contexts in which the therapeutic utility of PARPi can be expanded by targeting ALC1, whose inhibitors are currently in Phase I clinical trials.

## INTRODUCTION

High-grade serous ovarian cancer (HGSOCs) is the gynecologic malignancy with the highest case-fatality rate(*1*). Whole genome sequencing of HGSOCs revealed that while approximately 50% of these tumors have evidence of homologous recombination (HR) deficiency, another 20% are HR-proficient (HRP) and associated with amplification of an oncogene, *CCNE1* (cyclinE1-amplified)(*2*). The treatment regimen for HR-deficient (HRD) serous ovarian cancers includes Poly (ADP-ribose) polymerase inhibitors (PARPi), both as a frontline and maintenance therapy for platinum-sensitive tumors(*3*). However, major challenges associated with clinical success of PARPi include inevitable emergence of resistance and development of hematological disorders(*4*, *5*). Cyclin E1-amplified HGSOCs, on the other hand, display resistance to both platinum and PARPi-based therapies(*6*). Currently, there are no FDA-approved approaches to reduce effective PARPi dosage for alleviating toxicity or to make this targeted therapy effective in the context of cyclin E1-amplified serous ovarian cancers.

The potency of PARPi in killing HRD cells emanates from their ability to trap PARP1 and 2 enzymes on the chromatin(*7*, *8*). Trapped PARP1 and 2 enzymes impede fork progression and result in replication-coupled gaps and breaks, which increase the reliance on HR proteins for repair and survival. Hence, trapping potency of various PARPi directly correlates with cell cytotoxicity. In cells, talazoparib is the most potent PARP1/2 trapper and induces the most cytotoxicity, followed by niraparib. Olaparib and rucaparib share similar PARP1/2 trapping potency, albeit weaker than talazoparib and niraparib. Compared to these PARPi, veliparib is a weaker PARP1/2 trapper and hence it displays greatly reduced potency in cell killing(*7*, *8*). In concordance with higher trapping potential leading to cytotoxic side effects, incidences of anemia and cytopenia are more common with talazoparib and niraparib. In contrast, veliparib, despite its reduced toxicity and lack of susceptibility to multi-drug efflux pumps, has not shown meaningful efficacy in several clinical trials(*9*). Currently, niraparib, olaparib and rucaparib are approved as frontline and maintenance therapies for platinum-sensitive HGSOCs. To address the limitations of PARPi, there are ongoing efforts to design combinatorial strategies. However, it is unclear if different PARPi will display similar potency in killing cancer cells when used in combination with other agents.

We previously pursued a CRISPR screen to identify factors that can enhance PARPi sensitivity in UWB1.289, a *BRCA1*-mutant HGSOC cancer cell line(*10*, *11*). One of the top candidates that emerged from the screen was a nucleosome sliding enzyme called Amplified in Liver Cancer 1 (ALC1). More recently, we demonstrated that ALC1 mediated chromatin remodeling is specifically required for the repair of chromatin-buried abasic sites(*12*), an intermediate formed during base excision repair or generated by the spontaneous loss of a nucleobase from single-strand DNA(*13*). ALC1 is dispensable for responses to many other genotoxins(*12*). Furthermore, ALC1 knockout mice are viable and do not display a compromise in fitness or cancer predisposition(*11*). These unique features have propelled the entry of ALC1 inhibitors into a Phase I clinical trial (NCT06525298). However, the contexts in which ALC1 targeting will be most fruitful to enhance PARPi sensitivity remain unclear.

Here, by using a panel of established cancer lines and patient-derived cells we demonstrate that ALC1 loss enhances sensitivity to various PARPi across multiple HGSOC contexts, which include platinum-sensitive *BRCA1/2*-mutant, platinum-resistant *BRCA1/2*-mutant, as well as cyclin E1-amplified settings. We also highlight the safety associated with ALC1 targeting by uncovering that loss of this chromatin remodeler has minimal to no impact on genome stability and PARPi response in *BRCA*-WT and *BRCA*-heterozygous non-cancerous fallopian tube cell lines. Finally, guided by clinical responses to PARPi in ovarian cancer patients, we uncover that ALC1 targeting is likely to be most effective in tumors with high levels of endogenous replication stress, which can be quantified using phosphoT21-replication protein A2 (RPA2).

Together, this study suggests that targeting ALC1 can be a safe and clinically actionable approach to significantly improve PARPi efficacy in a wide spectrum of HGSOCs.

## RESULTS

### ALC1 loss enhances sensitivity to various PARPi in multiple BRCA1/2-mutant HGSOC cell lines

We have previously shown that ALC1 loss enhances olaparib sensitivity of UWB1.289, a *BRCA1*-mutant ovarian cancer cell line(*10*). We first examined whether this sensitivity could be extended to other PARPi. To do so, we depleted ALC1 using a previously established SaCas9-sgRNA strategy(*10*) and assessed sensitivity to olaparib, rucaparib, niraparib and veliparib. Consistent with our previous findings, ALC1 depletion significantly enhanced sensitivity of UWB1.289 cells to olaparib. In addition, *ALC1*-deficient UWB1.289 cells were more sensitive to rucaparib and veliparib as compared to their parental counterparts. Enhancement in sensitivity to niraparib upon ALC1 loss was relatively modest. A similar drug sensitivity profile upon ALC1 loss was also observed in JHOS4, another *BRCA1*-mutant HGSOC line (**Fig. 1a, b**). Importantly, we could recapitulate the enhanced sensitivity to olaparib and rucaparib upon ALC1 loss in cells derived from the ovarian tumor of a patient with a germline *BRCA*1 mutation (AOCS14)(*14*)(**Fig. 1c**) or with *BRCA1* promoter methylation (AOCS11.2)(*14*) (**Supp. Fig. 1**). Together, these findings suggest that targeting ALC1 can significantly enhance the sensitivity of *BRCA1-*mutant HGSOC cells to various PARPi.

**Figure 1.**
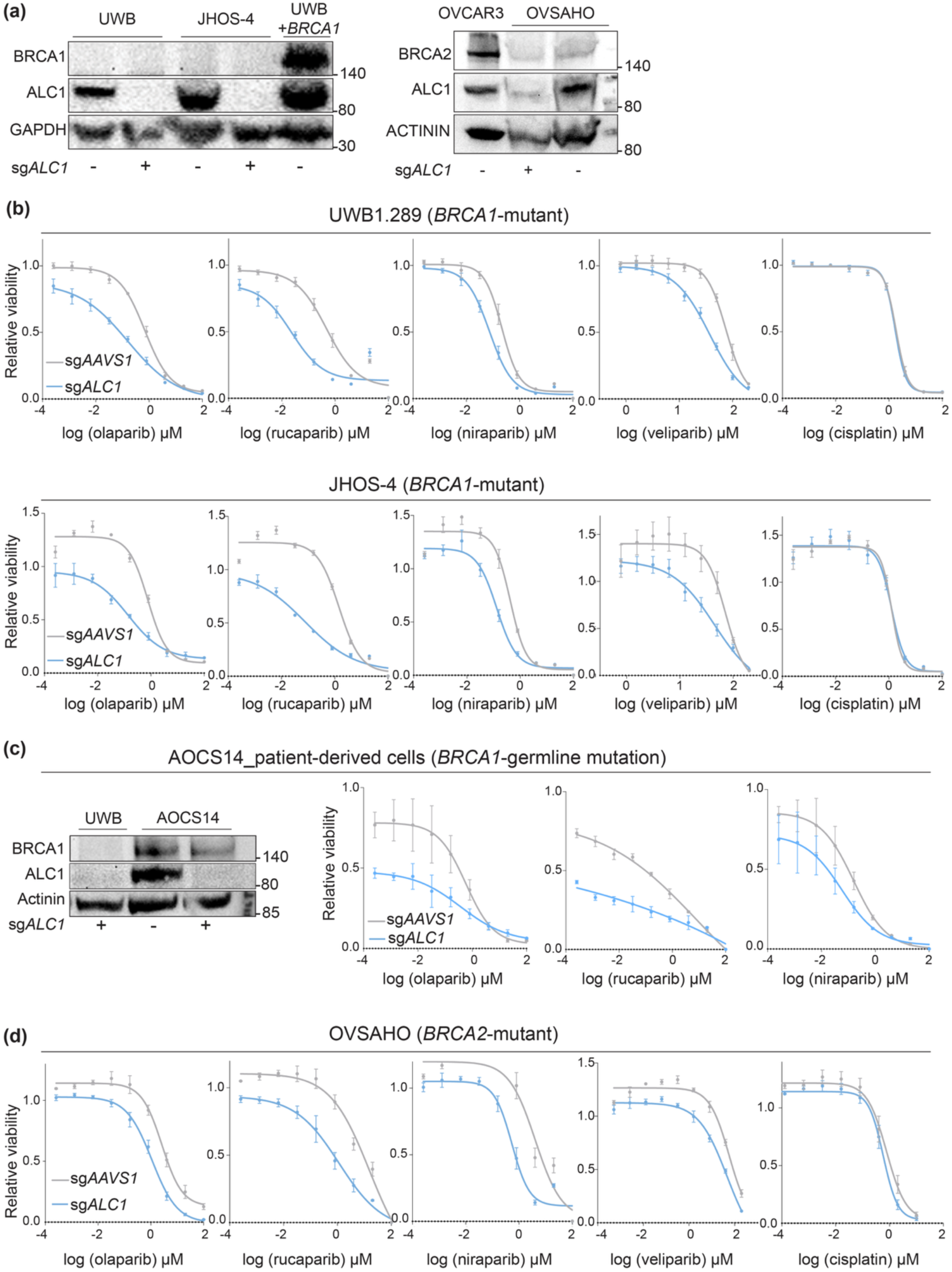
Loss of ALC1 enhances PARPi sensitivity across *BRCA1/2*-mutant ovarian cancer cells. **(a)** Immunoblot showing ALC1 depletion levels in indicated cell lines. **(b)** Sensitivities of the indicated *BRCA1*-mutated cell lines to various PARPi and Cisplatin using the CellTiter-Glo assay; n = 3 biologically independent experiments. Data are presented as mean ± s.e.m. **(c)** Immunoblot (left) showing ALC1 depletion levels and PARPi sensitivity (right) of the patient-derived cells with a *BRCA1* germline mutation quantified using the CellTiter-Glo assay; n = 3 biologically independent experiments. Data are presented as mean ± s.e.m. **(d)** Sensitivities of the indicated OVSAHO cell line to various PARPi and Cisplatin using the CellTiter-Glo assay; n = 3 biologically independent experiments. Data are presented as mean ± s.e.m.

We next examined the impact of ALC1 depletion on PARPi responses in the *BRCA2*-mutant HGSOC cell line, OVSAHO. ALC1 loss in OVSAHO cells enhanced sensitivity to olaparib, rucaparib and niraparib. Minimal enhancement in veliparib sensitivity was observed in *ALC1*-deficient OVSAHO cells (**Fig. 1a, d**). In both *BRCA1/2*-mutant settings, ALC1 loss showed stronger sensitization to rucaparib and olaparib, suggesting that the intermediate trapping potential of these PARPi may make them more favorable for combinatorial therapies^6^.

We also tested the impact of ALC1 loss on sensitivity to cisplatin, a platinum-based therapy. Notably, ALC1 loss did not enhance sensitivity to cisplatin in either *BRCA1* or *BRCA2*-deficient cells (**Fig. 1a, b**). Cisplatin-induced double-strand break (DSB) repair in HRD settings can be performed by Pol theta catalyzed microhomology-mediated end joining (MMEJ)(*15*, *16*). Our data showing the dispensability of ALC1 in cisplatin response, both in *BRCA1* and *2*-mutant settings, suggests that this chromatin remodeler likely does not contribute to MMEJ. ALC1 dispensability for MMEJ highlights that this chromatin remodeler has a different mechanism of action compared to Polymerase Theta, whose inhibitors are currently in clinical trials for treatment of multiple forms of HRD-ovarian cancers(*17*).

### ALC1 loss sensitizes HR-proficient cyclin E1-amplified cells and xenograft models to PARPi

Approximately 20% of HGSOCs are associated with amplification of *CCNE1*(*2*). Notably, *CCNE1* amplification and HR deficiency are almost entirely mutually exclusive. Individuals with cyclin E1-amplified ovarian cancers display poor prognosis and resistance to platinum-and PARPi-based treatments because of their proficiency in HR(*18*, *19*). We first examined whether HR-proficiency was retained in cell line models of cyclin E1-high (amplification or gain in *CCNE1*) HGSOCs. To test this, we used two cell lines, OVCAR3 and OVCAR4, which exhibit amplification in *CCNE1*, as well as OVCAR8, which shows a gain in *CCNE1* copy number(*20*). Upon challenging with olaparib, both OVCAR3 and 4 cells can form RAD51 foci comparable to *BRCA1*-addback UWB1.289 cells, establishing their HRP status (**Supp. Fig. 2**) OVCAR8 cells were also proficient in forming RAD51foci albeit at a level lower than OVCAR3 and 4. This could perhaps be attributed to the heterozygous methylation status of the *BRCA1* promoter in OVCAR8 cells(*21*).

Increased levels of cyclin E1 have been shown to induce high levels of replication stress, which results in the accumulation of single-strand DNA(*22*, *23*). Nucleobases in single-stranded DNA can be spontaneously lost resulting in the formation of abasic sites, lesions that underlie PARPi hypersensitivity in ALC1-deficient cells(*12*). We therefore reasoned that increased replication stress in cyclin E1-high cancer cells may render them sensitive to PARPi treatment upon ALC1 depletion. Consistent with this hypothesis, we observed that ALC1 depletion significantly enhanced olaparib and rucaparib sensitivity in OVCAR3, OVCAR4 and OVCAR8 cells (**Fig. 2a, b**). To test our hypothesis in a clinically relevant setting, we obtained patient cells from ovarian tumors with amplification (AOCS40) and gain (AOCS30)(*14*) in *CCNE1*(**Fig. 2c**). Depletion of ALC1 in both *CCNE1*-amplified and gain cells conferred enhanced sensitivity to olaparib and rucaparib (**Fig. 2d)**. In line with our prior observations, the impact of ALC1 loss on niraparib sensitivity was modest in cyclin E1-high cells (**Supp. Fig. 3**). Furthermore, ALC1 loss did not impact sensitivity to cisplatin, supporting its dispensability for HR repair (**Supp. Fig. 3**). Together, these observations demonstrate that targeting ALC1 can be an effective therapeutic approach to sensitize chemo-resistant cyclin E1-high ovarian cancers to olaparib and rucaparib.

**Figure 2.**
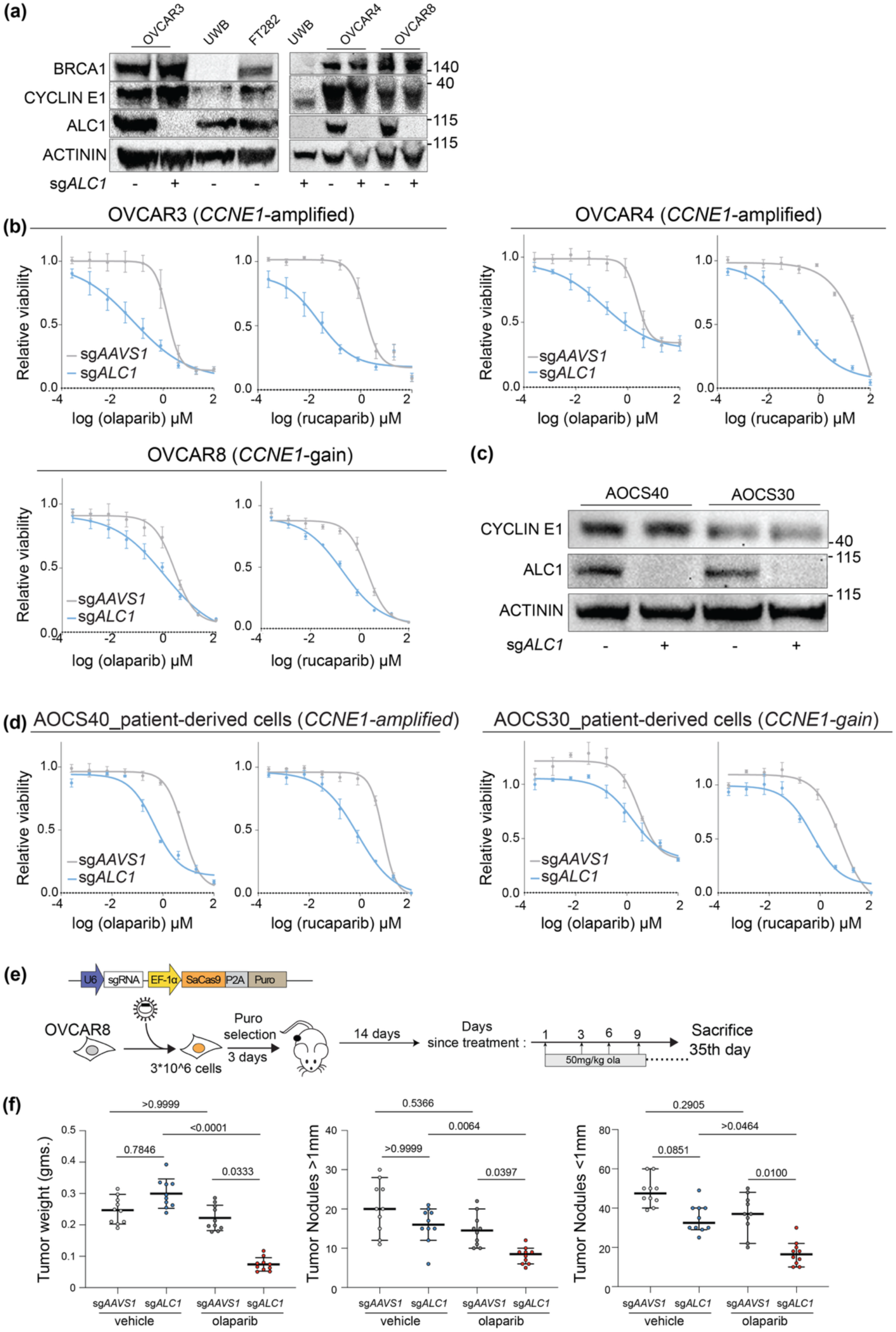
ALC1 loss enhances PARPi sensitivity cyclin E1-high ovarian cancer cells and xenograft models. **(a)** Immunoblot showing ALC1 depletion levels in indicated cyclin E1-high cancer cell lines. **(b)** Sensitivities of indicated cyclin E1-high cancer cell lines to various PARPi and Cisplatin using the Resazurin assay; n = 3 biologically independent experiments. Data are presented as mean ± s.e.m. **(c)** Immunoblot showing ALC1 depletion levels in indicated cyclin E1-high cells derived from ovarian tumors. **(b)** Sensitivities of indicated cyclin E1-high patient-derived cells to various PARPi and Cisplatin using the CellTiter-Glo assay; n = 3 biologically independent experiments. Data are presented as mean ± s.e.m. **(e)** Schematic for the xenograft experiment. **(f)** Assessment of tumor weight and nodules in indicated OVCAR8-derived xenografts. Mice were scarified at 35th day after initiation of PARPi treatment and indicated parameters were quantified. N=10/mice per group. P-value: Kruskal-Wallis test.

We next examined how ALC1 loss impacted PARPi response in a cyclin E1-high cell xenograft model. To do so, we utilized OVCAR8 cells, since they have been established to form ovarian tumors in mice(*24*). OVCAR8 cells were engineered to stably express sgRNAs targeting either *ALC1* or the safe locus *AAVS1* as a negative control. These cells were intraperitoneally injected into nude mice. At 14 days, two mice from each of the sg*AAVS1* and sg*ALC1*groups were sacrificed and analyzed to ensure development of ovarian tumor nodules. Starting on the 14th day after tumor injection, 10 mice from each of the sg*AAVS1* and sg*ALC1* groups were administered 50mg/kg of olaparib via oral gavage every third day, while another group was administered vehicle control. On the 35th day after initiation of PARPi treatment, mice were sacrificed to measure tumor weight and nodules (**Fig. 2e**). Consistent with the HRP status of OVCAR8, olaparib treatment in sg*AAVS1* group did not have an impact on tumor growth. In contrast, olaparib-treated sg*ALC1* cells showed significantly reduced tumor growth and tumor nodules (**Fig. 2f**). Collectively, our cell-based and xenograft studies suggest that targeting ALC1 can expand the utility of PARPi for the treatment of cyclin E1-high serous ovarian cancers, which currently have limited to no therapeutic options.

### ALC1 loss restores PARPi sensitivity across clinically relevant models of chemo-resistant ovarian cancers

Given that ALC1 loss can significantly enhance PARPi sensitivity in HR-proficient cyclin E1-high ovarian cancer cells, we next examined if this impact is also observed in chemoresistance models of *BRCA*-mutant ovarian cancers. *BRCA1*-mutant ovarian cancer cells eventually develop chemoresistance when exposed to PARPi for a prolonged period. Mechanistically, this is due to rewiring of ATR signaling, which enables RAD51 loading at DNA breaks and reversed forks independent of BRCA1 protein(*25*). This has been the key rationale for multiple ongoing clinical trials for ovarian cancers involving a combination of ATR inhibitor (ATRi) and PARPi. While the results of combining ATRi and PARPi have been promising across multiple clinical trials, the adverse side effects of cytopenia and severe anemia have limited its clinical translation(*26*). We examined whether ALC1 depletion might counteract PARPi resistance arising from the ATR-dependent repair compensating for the compromised BRCA1 activity. To do so, we procured two UWB1.289 cell lines, SYr12 and SYr14, that have acquired PARPi resistance due to increased reliance on ATR-mediated RAD51 loading at DNA double-strand breaks and reversed forks(*25*) (**Supp. Fig. 4a**). Notably, ALC1 loss imparts sensitivity to olaparib and rucaparib in both SYr12 and SYr14 cell lines. These data highlight that targeting ALC1 can be a viable approach to overcome PARPi resistance in *BRCA1*-mutant ovarian cancers that have bypassed the functional requirement of BRCA1 by increasing reliance on ATR activity (**Fig. 3a, b**).

**Figure 3.**
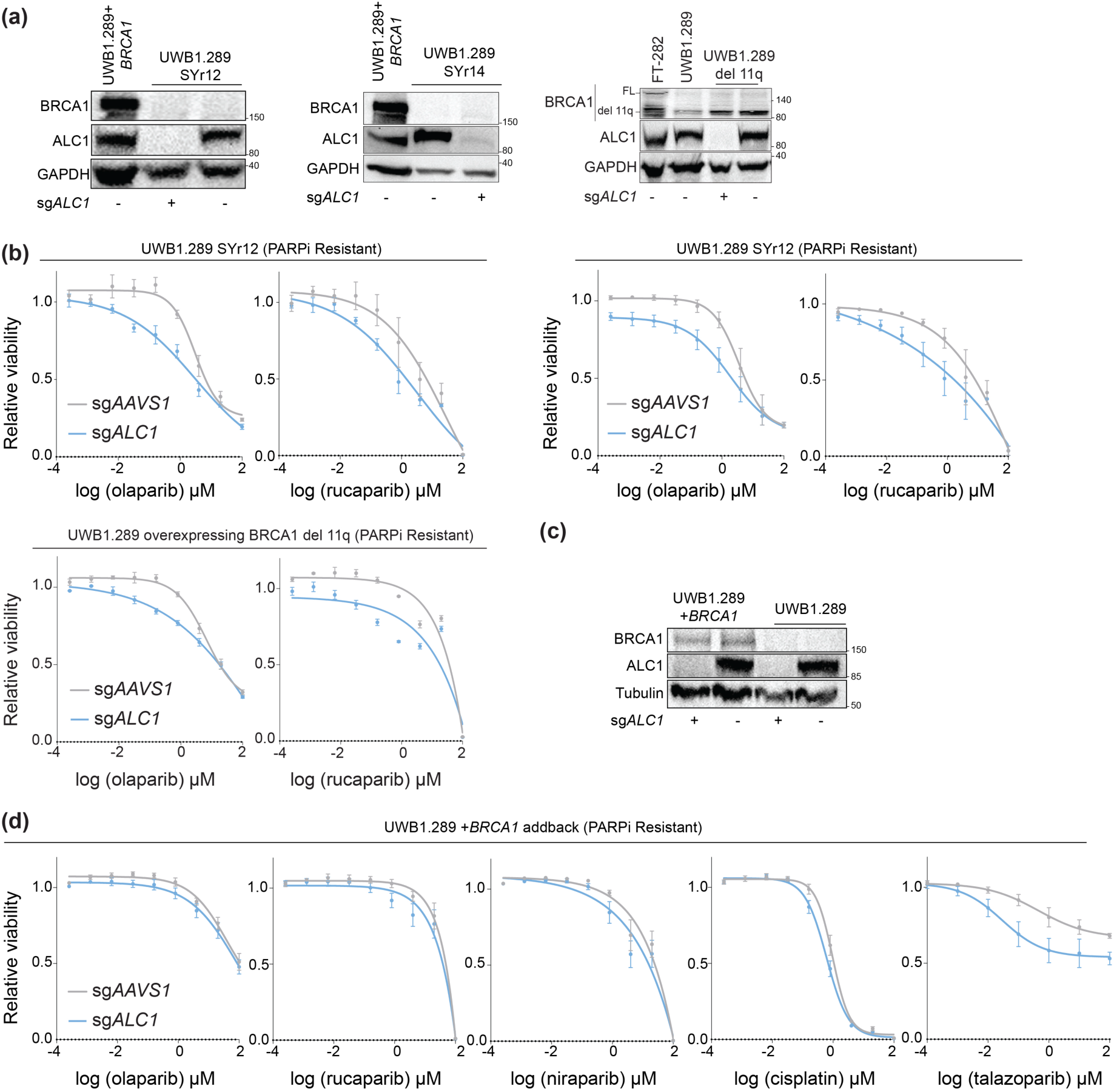
ALC1 loss restores PARPi sensitivity in ovarian cancer cell lines with partially restored homologous recombination. **(a)** Immunoblot showing BRCA1 and ALC1 depletion levels in indicated cell lines. **(b)** Sensitivities of indicated cells to various PARPi and Cisplatin using the CellTiter-Glo assay; n = 3 biologically independent experiments. Data are presented as mean ± s.e.m.. **(c)** Immunoblot showing BRCA1 and ALC1 depletion levels in indicated UWB1.289 cell lines. **(d)** Sensitivities of indicated UWB1.289+BRCA1 add-back cells to various PARPi and Cisplatin using the CellTiter-Glo assay; n = 3 biologically independent experiments. Data are presented as mean ± s.e.m..

Many pathogenic mutations in *BRCA1* do not abrogate the expression of the entire protein but instead result in overexpression of various hypomorphs. Overexpression of BRCA1 hypomorphs is yet another commonly reported mechanism of PARPi resistance in ovarian cancer(*27–31*) (**Supp. Fig. 4b**). To examine if ALC1 loss can restore PARPi sensitivity in ovarian cancer cells that have acquired resistance due to overexpression of hypomorphs, we procured UWB1.289 cells that are PARPi resistant due to overexpression of the del l1q isoform of BRCA1(*30*). This isoform is derived by alternative splicing wherein most nucleotides in exon 11 (c.788-4096) are lost. We observe that ALC1 loss potentiates olaparib and rucaparib sensitivity in del11q overexpressing resistant models (**Fig. 3a, b**). Finally, we explored the resistant setting wherein full length *BRCA1* is exogenously overexpressed in the UWB1.289 cell line(*32*). Consistent with the dispensability of ALC1 in HR, depletion of this remodeler in cells overexpressing fully functional BRCA1 protein did not enhance sensitivity to olaparib, rucaparib, niraparib or cisplatin. However, we observe that ALC1 loss can increase talazoparib sensitivity in cells overexpressing BRCA1 WT protein (**Fig. 3c, d**). Together, our studies suggest that while targeting ALC1 can be a viable approach to restore PARPi sensitivity in tumors that have partially restored HR, its therapeutic utility in HGSOCs with BRCA1 overexpression could be limited. Of note, impact of ALC1 on PARPi response is more profound in HRD and cyclin E1-high settings compared to scenarios of acquired resistance. These observations suggest that combining ALC1 targeting with PARPi in an upfront setting is likely to be more efficacious compared to when ovarian tumors have acquired chemoresistance.

### ALC1 loss does not impact PARPi responses or genome stability in *BRCA*-WT and heterozygous fallopian tube models

The minimal impact of ALC1 in *BRCA1*-proficient UWB1.289 cells suggested that targeting this remodeler may have minimal impact on normal healthy cells. To test this, we used a panel of cell lines derived from normal fallopian tube (FT) epithelium, the tissue where the majority of HGSOCs originate(*33*, *34*). The FT cell line panel included hTERT-immortalized FT190, FT194 and FT282. Remarkably, ALC1 loss did not enhance sensitivity to either platinum or any of the clinically approved PARPi (**Fig. 4a, b**). We next extended our studies in the context of non-cancerous *BRCA1*-heterozygous FT epithelial cells. This scenario represents normal healthy cells in individuals who are born with a pathogenic copy of the *BRCA1* or *2* gene and hence are heterozygous for the mutation at the organismal level(*35*). To do so, we used CRISPR/Cas9 to generate two different FT282 clones with *BRCA1* heterozygous genotype. We confirmed the heterozygous status of the cell line using next-generation sequencing and western blotting (**Supp. Fig. 5**). ALC1 loss had minimal to no impact on PARPi or platinum responses in *BRCA1*-heterozygous FT282 cells (**Fig. 5a, b**), reinforcing the potential safety associated with targeting this chromatin remodeler for therapeutic purposes.

**Figure 4.**
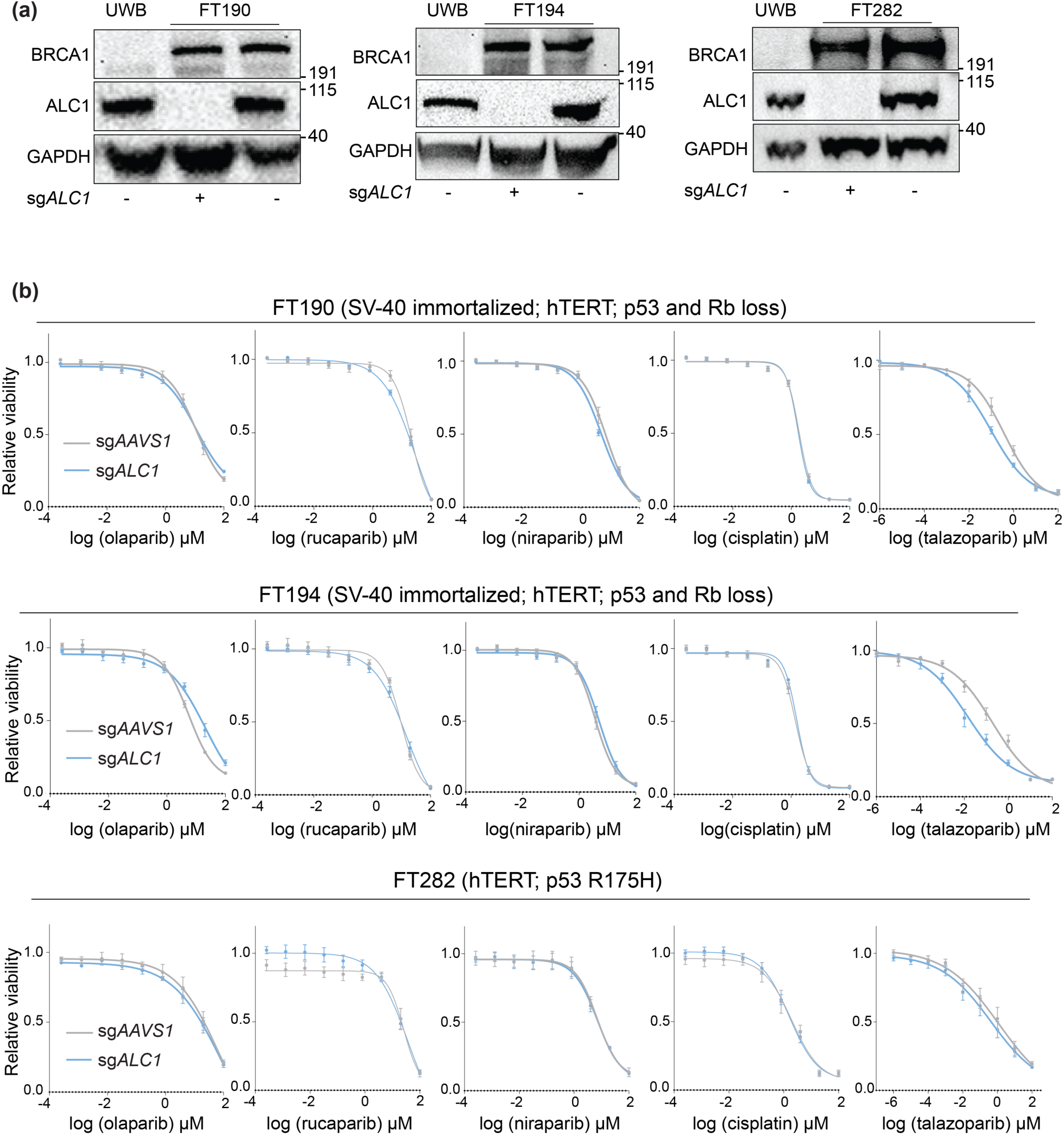
ALC1 loss does not enhances PARPi or cisplatin sensitivity in *BRCA1/2*-proficient non-cancerous immortalized fallopian tube (FT) cell lines. **(a)** Immunoblot showing ALC1 depletion levels in indicated FT-derived cell lines. **(b)** Sensitivities of the indicated FT-derived cell lines to various PARPi and Cisplatin using the Resazurin assay; n = 3 biologically independent experiments. Data are presented as mean ± s.e.m.

**Figure 5.**
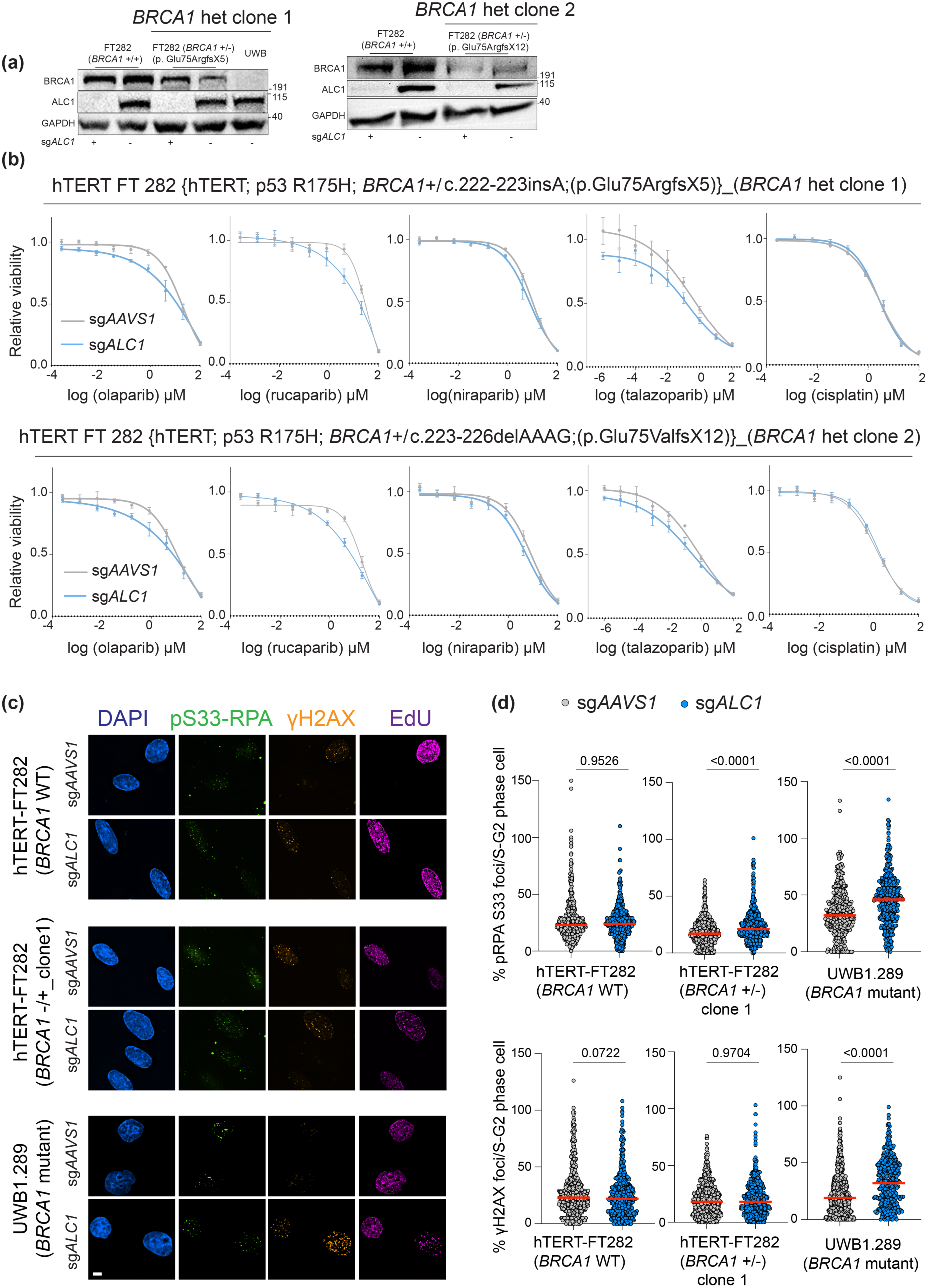
Loss of ALC1 does not enhance PARPi sensitivity or genome instability in *BRCA1*-heterozygous fallopian tube cell lines. **(a)** Immunoblot showing ALC1 depletion levels in indicated fallopian tube-derived *BRCA1*-het clones. **(b)** Sensitivities of the fallopian tube-derived *BRCA1*-het clones to various PARPi and Cisplatin using the Resazurin assay; n = 3 biologically independent experiments. Data are presented as mean ± s.e.m. **(c)** Representative images of pS33-RPA and gH2AX in indicated cell lines. Brightness and contrast of the images are constant between sg*AAVS1* and sg*ALC1* pair of a given cell line but are different between different cell types. Scale bar 10 micron. EdU: 5-Ethynyl-2 deoxyuridine. **(d)** Quantification of pS33-RPA (top) and gH2AX (bottom) foci per S/G2 cells. n= 3 biologically independent experiments. Median is indicated in red. P-value: Mann Whitney. For examining cell cycle stages, DAPI intensities of EdU positive cells (S-phase cells) was averaged. Any EdU negative cells with DAPI intensity less and more than the average S phase was referred to as G1 and G2 respectively.

To further establish that ALC1 targeting is likely to have minimal impact on normal non-cancerous cells, we assessed how loss of this remodeler impacts genome stability in *BRCA1*-WT FT282, *BRCA1*-heterozygous FT282 and *BRCA1*-mutant UWB1.289 cells. We determined levels of replication stress and replication-coupled double-strand breaks by quantifying pS33-RPA2 and γH2AX respectively. ALC1 loss did not enhance either pS33-RPA or γH2AX levels in *BRCA1*-WT FT282 cells. In *BRCA1*-heterozygous FT282 cells, ALC1-deficiency showed a modest 1.2-fold increase in pS33-RPA2 with no concomitant increase in γH2AX. The modest enhancement in pS33-RPA could be due to the inability of *BRCA*-heterozygous cells to resolve replication stress(*36*, *37*), which can be generated upon ALC1 loss. Unresolved replication stress eventually results in DNA double-strand breaks. We reasoned that the proficiency of *ALC1*-deficient *BRCA1*-heterozygous cells in HR repair can eventually resolve any genome instability that may arise due to the modest increase in replication stress. Consistent with this hypothesis, we do not observe an increase in double-strand breaks in ALC1-deficient *BRCA1*-heterozygous cells compared to their ALC1-proficient counterparts. Unlike in *BRCA1*-WT and *BRCA1*-heterozygous FT282 cells, ALC1 loss in *BRCA1*-mutant UWB1.289 cells resulted in a 1.4- and 1.7-fold increase in pS33-RPA or γH2AX respectively (**Fig. 5c, d**). Together, these observations strengthen the validity of ALC1 as a safe target while concomitantly hypersensitizing multiple forms of HGSOCs to PARPi.

### Enhanced PARPi sensitivity upon ALC1 loss can be predicted by the endogenous levels of replication stress

We next extended our cell-based findings to clinical samples to uncover how ALC1 levels correlate with PARPi response in ovarian tumors. We first examined publicly available The Cancer Genome Atlas (TCGA) data for 602 HGSOCs deposited in cBioPortal to assess for genomic alterations in ALC1(*38*). Approximately 7% of cases were associated with amplification in *ALC1* (**Fig. 6a**), suggesting an increased expression of this gene in ovarian cancer. In concordance, assessment of ALC1 protein levels using immunohistochemistry analysis of tissue-microarray (TMA) derived from individuals with HGSOCs revealed a significantly higher protein expression in primary serous ovarian tumors (n=142) as compared to benign tissue samples (n=51) (**Fig. 6b**). Higher levels of ALC1 in tumors therefore provide the opportunity to target this protein for therapeutic utility. We next examined previously published proteomics data from HRD HGSOC tumors to assess the correlation between ALC1 protein levels and estimated time individuals were on PARPi treatment (**Supp. Table 1**)(*39*). We observe that low ALC1 protein levels trend with longer time on PARPi (**Fig. 6c)**, revealing that reduced expression of this chromatin remodeler likely results in a more durable response in HRD-ovarian cancer patients. Of note, this inverse correlation is not owing to a decrease in ALC1 protein levels over time during treatment, ruling out the possibility for a negative selective pressure (**Supp. Fig. 6**).

**Figure 6.**
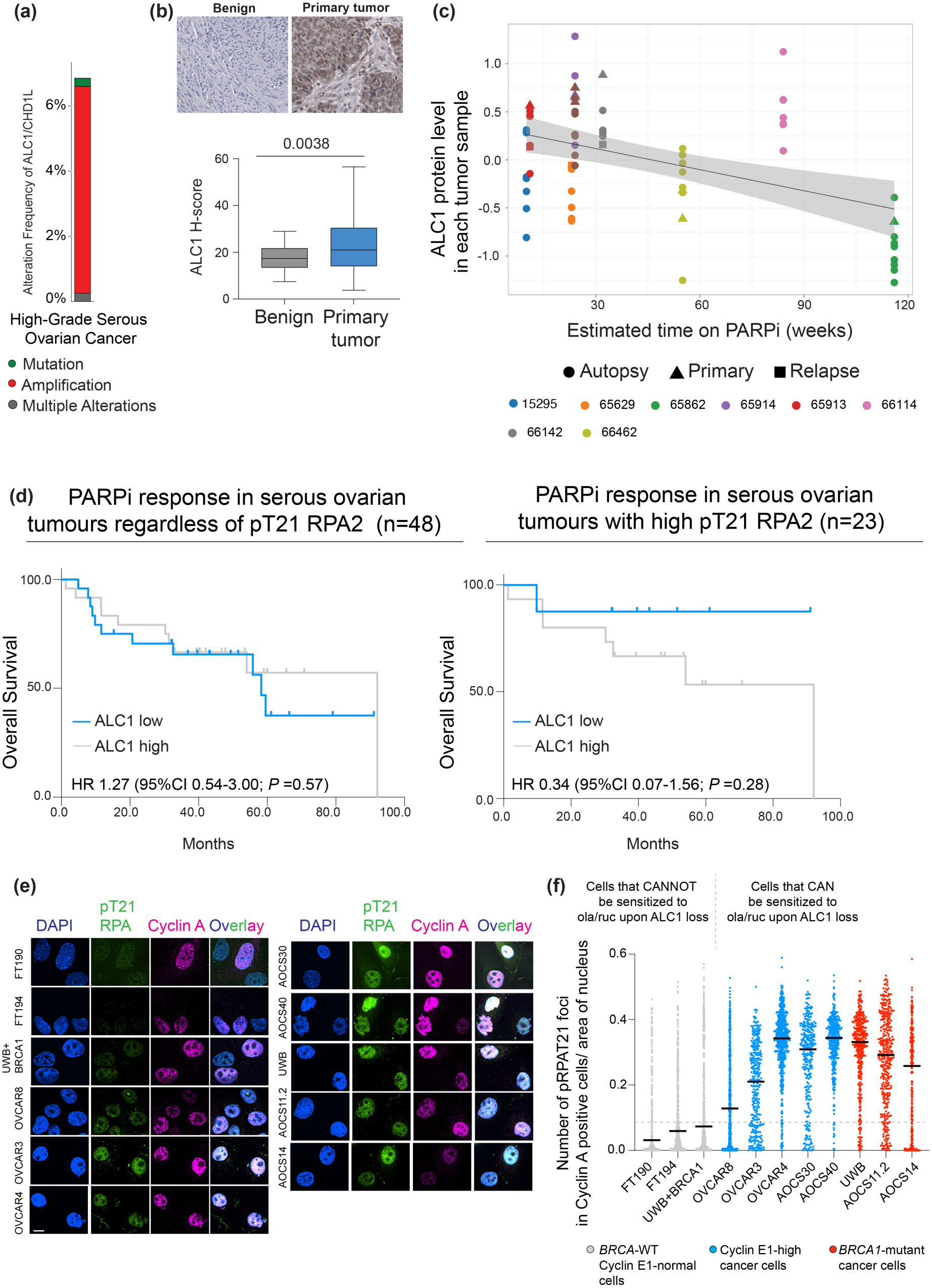
ALC1 targeting exploits cancer cell intrinsic replication stress. **a.** Bar graph taken from cBioPortal showing alteration frequency in ALC1 in HGSOCs. **b.** Representative image showing ALC1 protein expression in indicated tissues (top) and box-and-whisker plots showing quantification of ALC1 protein expression levels per H-scores in normal ovarian tissue (n=51 sites) and primary tumor (n=142 different sites) sampled from 78 patients. P-value is derived from Mann-Whitney test. **c.** Graph showing correlation with ALC1 protein levels and estimated time on PARPi. Due to the small size of the cohort, no statistical analysis was performed. The case number corresponds to the IDs from Burdett *et al*, 2023. **d.** Kaplan-Meier curves evaluating overall survival with PARPi in individuals without (left) or with (right) pT21 RPA2-high ovarian tumors. P-value is derived from Logrank (Mantel-Cox) test. **e.** Representative images showing levels of pT21 RPA levels in Cyclin A positive cells. Scale bar: 10 microns. **f.** Quantification of pT21 RPA levels in Cyclin A positive cells normalized by nuclear size in indicated cell lines. Median is indicated.

Given our cell-based studies suggest that ALC1 targeting can have an impact beyond *BRCA1/2*-mutant settings, we evaluated expression of this remodeler in a TMA derived from an independent cohort of PARPi treated patients (n=48), which included both HRD and HRP samples^40^. Of these, 33 and 15 patients had International Federation of Gynecology and Obstetrics (FIGO) stage III and IV HGSOC respectively. 19 patients had *BRCA* mutations. 28 patients received PARPi in the frontline maintenance setting and 20 at the time of recurrence. The median follow-up was 56.1 ± 25.2 months (13.9-146.4). Samples were classified as ALC1-High (≥23% ALC1-positive staining) or ALC1-Low (<23%), with the cutoff determined by systematic evaluation of all observed ALC1 values and selection of the threshold that minimized the P value^40^. Since tumor biopsies were obtained at the time of diagnosis and patients received PARPi in both the frontline maintenance and recurrent settings, progression free survival (PFS) was not an optimal endpoint. Hence, we prioritized overall survival (OS) after PARPi treatment as a measure of response in this heterogeneous treatment context. No significant association was observed between ALC1 expression and OS in the large cohort (**Fig. 6d, left**). Mechanistically, ALC1 targeting exploits cancer cell intrinsic HR deficiency or increased replication stress^12^. We therefore hypothesized that ALC1 expression would correlate with PARPi response in HRD tumors, as quantified by RAD51 foci, or in tumors with high levels of replication stress, as quantified by phosphorylation of T21 residue of RPA2(*40*). Due to limited numbers in the HRD cohort (n=9), correlation between ALC1 level and OS could not be evaluated in this set. However, in the HRP, high replication stress cohort (n=23), patients with ALC1-Low tumors had longer OS than those with ALC1-High tumors (undefined versus 92.1; hazard ratio 3.1, 95% CI 0.4-25.5, *P*=0.3) (**Fig. 6d, right**). Although the effect size was large, it did not reach statistical significance. This is likely due to limited events, as 87.5% of cases in the ALC1-Low, HRP, pRPA-high group were censored because recurrence had not yet occurred at the time of analysis. Together, these patient-derived data highlight that ALC1 targeting is likely to benefit any tumor with high levels of intrinsic replication stress.

To experimentally validate that high replication stress can indeed predict the efficacy of ALC1 targeting in enhancing PARPi response, we quantified endogenous levels of pT21 RPA2 in a panel of HRP and HRD cells. These included cell types that can or cannot be sensitized to PARPi upon ALC1 depletion. We co-stained for pT21 RPA2 and S/G2 marker, Cyclin A, to alleviate any differences that may confound our findings due to differences in cell cycle distribution across various lines. Furthermore, the number of pT21 RPA foci was normalized to the area of the nucleus to account for the difference in the size of various cell types used for the analysis. We observe that any cell type with a median greater than 0.1 pT21 RPA foci/nuclear size is sensitive to olaparib and rucaparib upon ALC1 depletion (**Fig. 6e, f)**. Together our data strongly supports that ALC1 targeting can efficiently and safely expand the therapeutic efficacy of PARPi in tumors with high endogenous levels of replication stress. Our findings also warrant that pT21 RPA2 can be investigated further as a functional biomarker to predict PARPi response upon ALC1 targeting.

## DISCUSSION

Here, we define the clinical context in which targeting ALC1 can effectively overcome the clinical limitations of PARPi in the regimen of HGSOCs. We observe a profound impact of ALC1 depletion on PARPi responses in ovarian cancer cells with *BRCA1/2*-mutation (**Fig. 1**), with high cyclin E1-levels, (**Fig. 2**) and with partial restoration of HR (**Fig. 3**). Our data suggests that targeting ALC1 can permit low PARPi dosage for an effective response, thereby potentially reducing therapy-associated side effects. Moreover, the expedited PARPi-mediated killing of cancer cells upon ALC1(*10*)also warrants that targeting this remodeler will likely be more beneficial in a frontline setting to mitigate the acquisition of resistance. In contrast, ALC1 targeting might likely not be beneficial in tumors with fully restored HR and low levels of endogenous replication stress (**Fig. 3d, 6e, f)**.

Many factors previously identified to enhance PARPi sensitivity, such as XRCC1, Fanconi anemia protein (FA), RAD51 and RAD54 paralogs, do so both in cancer and non-cancer cell lines(*41*). This is in part due to the roles of these proteins in either single-strand or HR repair and thus a compromise in the front-line repair mechanisms, resulting in enhanced sensitivity both in cancer and non-cancer cells. Similarly, while inhibitors of checkpoint kinase ATR or Chk1 synergize with PARPi in killing HGSOCs, their critical roles in DNA replication impose the risk of unwanted pathological phenotypes in normal healthy cells(*42*). In contrast, we demonstrate that ALC1 targeting has minimal to no impact on genome stability and PARPi sensitivity of normal fallopian tube epithelial cells (**Fig. 4, 5**). These data uncover the unique potential of ALC1 targeting in significantly expanding the therapeutic index of PARPi.

This distinction in the impact of ALC1 on cancer versus non-cancer cells prompted us to examine the endogenous vulnerability that is exploited upon ALC1 loss for enhanced PARPi response. We recently reported that PARPi hypersensitivity upon ALC1 loss is reliant on the presence of abasic sites(*12*). Abasic sites can arise spontaneously in genome upon the loss of a nucleobase, a process which tends to be four times higher in the context of single-stranded DNA compared to its double-stranded counterpart(*43*). We therefore speculated that replication stress, which results in the generation of single-stranded DNA, can directly correlate with increased PARPi sensitivity upon ALC1 loss. Indeed, we observe that low ALC1 levels manifest in enhanced PARPi response only in settings with high levels of replication stress, as quantified by the pT21 RPA2 (**Fig. 6d-f**). This observation could explain why cyclin E1-high cells, which harbor elevated replication stress, can respond to PARPi upon ALC1 loss despite their HRP status. The simplicity of an immunofluorescence assay to quantify pT21 RPA2 levels also highlights the potential of this approach to be adapted as a biomarker for predicting PARPi response upon ALC1 targeting.

As per FDA mandate, PARPi are typically prescribed to individuals who show positive responses to platinum. However, our data showing that ALC1 selectively sensitizes to PARPi, but not platinum-based drugs suggest that are distinctions in determinants of the responses to the two therapies frequently used in the regimen of HGSOCs. This is further reinforced by our observations that ALC1 targeting can allow platinum-resistant tumors to be sensitized to PARPi but not cisplatin. Hence, it might be worthwhile to reconsider the restriction of PARPi to only platinum-responsive tumors.

When implementing synergistic combinations, it is likely that not all PARPi will behave the same. Our studies simultaneously comparing the impact of available inhibitors that trap PARP1/2 enzymes reveal that moderate trappers, olaparib and rucaparib, are likely to be more favorable in combinatorial therapy. We also show that ALC1 targeting in *BRCA1*-mutant cancers can permit the utility of veliparib, which has lesser side effects compared to other clinically approved PARPi and also can be effective in situations where tumors have acquired resistance owing to activation of multi-drug efflux pumps(*44*). In contrast, in ovarian cancers that have acquired PARPi resistance due to overexpression of *BRCA1/2* protein, ALC1 targeting can enhance sensitivity to talazoparib. Hence, our studies warrant that in the context of combinatorial strategies, responses across PARPi can widely vary, and this should be an important clinical consideration.

Finally, we recognize that our studies are performed using ALC1 depletion and not using small molecule inhibitors of ALC1, which are not commercially available. However, our genetic studies and patient-derived data provide a comprehensive understanding of the relevant clinical avenues where pharmacological inhibitors of ALC1 will be meaningful.

## MATERIALS AND METHODS

### Immunohistochemistry (IHC) Staining

Tissue microarray studies were conducted on surgically resected high-grade serous ovarian cancer specimens from 78 patients diagnosed at the Department of Pathology at Washington University (St. Louis, MO, USA). All 5 tissue microarray slides were stained, imaged, and analyzed in parallel to minimize technical variability. Slides were pretreated for staining with a 45-minute bake step, and a 12-minute dewax step, followed by 20 minutes of antigen retrieval in citrate buffer at 95°C. Endogenous peroxidases were quenched in dilute hydrogen peroxide. Tissues underwent a 15-minute blocking step each with an Avidin D and Biotin blocking reagent (Vector Labs, SP-2001), followed by a 5-minute blocking step with Protein Block Serum-Free (Dako, X0909). Tissues were then incubated with Anti-CHD1L antibody (Sigma-Aldrich, HPA028670) overnight at 4°C. After washing off the primary antibody, tissues were incubated with the secondary antibody (Vector Labs, BA-1000) for 30 minutes at room temperature, followed by an incubation with horseradish peroxidase (Vector Labs, SA-5004) for 30 minutes at 37°C. Tissue-bound immunocomplexes were detected by incubating with diaminobenzidine (Dako, K3468) for 5 minutes at room temperature. Finally, tissue sections were counterstained with hematoxylin (BioCare Medical, CATHE-M). Stained tissue sections were imaged with an AxioScan.Z1 whole tissue scanner and ZEN v2 software (Carl Zeiss). Unbiased parallelized analysis was then performed with HALO v3 digital pathology (Indica Labs). Section images were annotated to include tissue regions of interest and exclude artifacts and debris. For the “Area Quantification v2” analysis module, stain colors and thresholds were set and measured across all sections to calculate the percentage of stains within the annotated regions of interest.

### Analysis of co-relation between ALC1 protein expression and PARPi response

Proteomic data was previously generated from laser micro dissected enriched tumor material from HGSOC patients with HRD disease(*39*). Of these cases, 9 patients had received PARPi treatment. A total of 65 tumor samples, collected at 3 timepoints (P= primary, R = relapse, A = autopsy), had proteomics data from these 9 patients (**Table 1**).

### Cell Culture

UWB1.289, UWB1,289+BRCA1, hTERT FT282, hTERT FT190, hTERT FT194, OVCAR3, OVCAR4, and OVCAR8 were procured from ATCC. OVSAHO cells were obtained from Sigma. UWB1.289 cells, including the derivative SYr12, SYr14, RR, and BRCA1-add back variants, were cultured in 1:1 Mammary Epithelial Cell Basal Medium (Thermo Fisher Scientific, M-171) and RPMI1640 with L-glutamine (Genesee Scientific, 25-506) supplemented with Mammary Epithelial Growth Supplement (Thermo Fisher Scientific, S0155), 10% fetal bovine serum (FBS) (Sigma, F0926), and 1x Penicillin-Streptomycin (P/S) (Genesee Scientific 25-512). OVSAHO, OVCAR8, and primary cell lines (AOCS) were cultured in RPMI1640 with L-glutamine (Genesee Scientific, 25-506) supplemented with 10% FBS and 1x P/S. OVCAR3 and OVCAR4 cells were cultured in RPMI1640 with L-glutamine (Genesee Scientific, 25-506) supplemented with 10% FBS, 1x P/S, and 0.5% insulin (Thermo Fisher Scientific, ITS-G 100X). hTERT FT190, hTERT 194, hTERT FT282, and FT282 BRCA1 heterozygous cells were cultured in DMEM/F-12 with L-glutamine (Sigma D8062) supplemented with 10% bovine calf serum (BCS) (Cytiva, SH30072.04) and 1x P/S. All AOCS cell lines were obtained from Australian Ovarian Cancer Study Group and have been previously described^14^.

### Lentivirus Transduction

HEK293T cells were used to produce lentiviruses. In a 6-well plate, HEK293T cells were transfected using Opti-MEM (Thermo Fisher Scientific, 31985070) solution containing 3750ng PAX2, 2500ng VSVG, 5250ng plasmid of interest, and 40uL of 1mg/mL polyethylenimine. 7 hours post-transfection, the media was changed. Lentivirus was collected 4 times over the course of 3 days, then stored at -80°C until time of use. Cells were transduced with lentivirus and 8ug/mL polybrene, followed by centrifugation at 600xg for 30 minutes to increase infection efficiency. 24 hours post-transduction, the media was changed. 48 hours post-transduction, a selection drug was added (Puromycin 2ug/mL or Blasticidin 5ug/mL).

### sgRNAs used in the study

The 2-in-1 SaCas9 system was used to deplete ALC1 for most of the cell viability studies. The two-vector SpCas9 system was used for depleting ALC1 and pol theta in the MMEJ reporter assay. These sgRNAs have previously been validated for target specificity. Nucleofection of guides with recombinant Cas9 protein was used to generate BRCA1 heterozygous FT282 cells. The sgRNAs used in the study are:

SaCas9-sg*ALC1*: 5’TCCACAACAAGAACACTCCAA 3’

SaCas9-sg*AAVS1*: 5’ GGTTCTGGGAGAGGGTAGCG 3’

SpCas9-sg*BRCA1*: 5’ TAAATCTCGTACTTTCTTGT 3’

SpCas9-sg*ALC1*: 5’ GTCGCCTGCATATGTTACAC 3’

SpCas9-sg*Pol theta:* 5’ ACTCCGACGCAGAAGATTCA 3’

SpCas9-sg*AAVS1:* 5’ GGGGCCACTAGGGACAGGAT 3’

### Immunoblotting

Cell pellets were lysed by incubation with a 5x volume of radioimmunoprecipitation assay (RIPA) buffer supplemented with 1mM dithiothreitol (DTT), 1x complete EDTA-free protease inhibitor cocktail (Sigma, 11873580001), 0.5uL benzonase (EMD Millipore, 70746-3), and 1mM phenylmethylsulfonyl fluoride (PMSF) for 20 minutes on ice. The cell suspension was then centrifuged at 4°C for 20 minutes to separate the supernatant from the cell debris. Bradford assay was used to quantify the protein concentration in the supernatant. 10-20ug of protein lysate was run on a Nupage 4-12% Tris protein gel (Thermo Fisher Scientific, NP0336) in 1x 3-(N-morpholino) propanesulfonic acid (MOPS) buffer. Protein was then transferred to a 0.2micron nitrocellulose membrane (Cytiva, 10600004) using a wet transfer at 400mA for 2 hours. Blocking was done in 5% milk in PBS with 0.2% Tween for 1 hour. Blots were incubated overnight with primary antibodies at 4°C.

The next day, blots were then washed 4 times with 0.2% Tween in PBS. Blots were next probed with HRP-conjugated secondary antibody at 1:3000 at room temperature for 1 hour, followed by 4 washes with 0.2% Tween in PBS. The Western lighting Plus ECL kit (Perkin Elmer, NEL10500) was used to develop the blot, followed by imaging on the iBright CL1500 Imaging system (Thermo Fisher Scientific, A44240). The details of primary and secondary antibodies are listed in **Supp. Table 2**.

### Cell Viability Assays

For all cell viability assays, sgRNA depletion of ALC1 or AAVS1 was completed 2 weeks prior to plating. For each cell type, 1000 cells in a volume of 100uL were plated in a 96-well plate in technical triplicates. UWB1.289, including SYr12, SYr14, del 11q, and BRCA1+ variants, JHOS-4, OVSAHO, and primary cells (AOCS) were plated in white 96-well plates (Alkali Scientific TPW90960). hTERT FT190, hTERT FT194, hTERT FT282 BRCA1 WT and heterozygous, OVCAR3, OVCAR4, and OVCAR8 cells were plated in clear 96-well plates. 24 hours after cells were plated, 100uL of 2x drug dilutions were added. For HR-proficient cell lines, viability was assessed 1 week after cells were plated. For HR-deficient cell lines, viability was assessed 1 week after drugs were added to the cells. Cell lines that were plated in clear 96-well plates were assessed using the Resazurin fluorescence assay. Accordingly, on the day of viability assessment, media was replaced with phenol red-free DMEM (Thermo Fisher Scientific, 31053028) supplemented with 10% bovine calf serum (BCS) (Cytiva, SH30072.04), 1x Penicillin Streptomycin, 2mM L-glutamine (Sigma D8062), and 10µg/ml resazurin (Sigma, 199303). Cells were incubated for 3-4 hours at 37°C in resazurin-containing media, followed by fluorescence measurement. Fluorescence was measured using the SoftMax Pro 6.4 software on the SpectraMax i3x microplate reader on 560nm excitation and 590nm emission settings. Cell lines that were plated in white 96-well plates were assessed using CellTiter Glo 2.0 Assay (Promega, PRG9242). Luminescence was measured on the aforementioned microplate reader on the luminescence setting. Fluorescence or luminescence of the drug-treated population was normalized to its respective untreated control, then survival curves were plotted. GraphPad Prism was used to fit the data to a curve via the following equation: y = min + (max – min) / 1 + 10^logEC50-x^.

### Immunofluorescence Staining

For each cell type, 25,000 cells were plated on poly-lysine-coated coverslips in a 24-well plate. Cells were given 48 hours to attach to the coverslips. For RAD51 foci formation experiments, cells were then treated for 24 hours with 10µM olaparib. 72 hours after plating, cells were pulse-labelled with 1ug/mL 5-Ethynyl-2’-deoxyuridine (EdU) for 20 minutes to label replicating cells. Following EdU labelling, cells were pre-extracted on ice for 5 minutes with ice-cold cytoskeleton buffer (10 mM Pipes pH 6.8, 100 mM NaCl, 300 mM sucrose, 3mM MgCl_2_, 1mM EGTA, 0.5% Triton X-100). For samples in which Cyclin A was used as the cell cycle marker, cells were not pulse-labelled or pre-extracted prior to fixation. Cells were fixed with 3% paraformaldehyde (PFA) for 15 minutes at room temperature. Four PBS washes were done to remove residual PFA following fixation. Cells were then permeabilized on ice for 5 minutes with 0.5% Triton-X in PBS, followed by blocking in 3% bovine serum albumin (BSA) dissolved in PBS. Incorporated EdU was labeled with Alexa-Fluor-647 Azide using the Click Reaction denoted in the manufacturer’s instructions (Fisher Scientific, 502108139). Cells were then washed once with the blocking buffer, 3 times with PBST, and once with PBS. Cells were fixed again with 3% PFA for 10 minutes at room temperature, then washed four times with PBST. For RAD51 and γH2AX staining, samples were incubated overnight at 4°C with rabbit-anti-RAD51 (ABCAM, ab63801) diluted 1:250 and mouse-anti-γH2AX (Sigma-Aldrich, 05-636) diluted 1:1000 in DMEM media with 0.02% sodium azide. For Cyclin A and pT21-RPA32 staining, samples were incubated overnight at 4°C with mouse-anti-Cyclin A (Santa Cruz, sc-271682) diluted 1:250 and rabbit-anti-pT21-RPA32 (ABCAM, AB61065) diluted 1:500 in DMEM media with 0.02% sodium azide. For γH2AX and pS33-RPA32 staining, cell samples were incubated for 1 hour at room temperature with mouse-anti-γH2AX (Sigma-Aldrich, 05-636) diluted 1:1000 and rabbit-anti-pS33-RPA32 (Thermo Fisher Scientific, A300-246A) diluted 1:500 in DMEM media with 0.02% sodium azide. Following four washes with PBST, samples labelled with anti-γH2AX and anti-RAD51 or anti-pS33-RPA32 were incubated for 1 hour at room temperature with Alexa-Fluor-555-conjugated anti-mouse (Thermo Fisher Scientific, A32773) and Alexa-Fluor-488-conjugated anti-rabbit (Thermo Fisher Scientific, A32731) secondary antibodies, each diluted 1:1000 in DMEM media with azide. Following four washes with PBST, samples labelled with anti-pT21-RPA32 and anti-Cyclin A were incubated for 1 hour at room temperature with Alexa-Fluor-647-conjugated anti-mouse (Thermo Fisher Scientific, A32787) and Alexa-Fluor-488-conjugated anti-rabbit (Thermo Fisher Scientific, A32731) secondary antibodies, each diluted 1:1000 in DMEM media with azide. Following four washes with PBST and four washes with MQ water, samples were incubated for 2 minutes at room temperature with 1μg/mL 4’,6-diamidino-2-phenylindole (DAPI) made in MQ water. Coverslips were mounted using Vectashield Vibrance Antifade Mounting Medium (Vector Labs, H-1700). Images were acquired using a Nikon Ti-2 microscope with a 60x objective. Z-stacks were imaged every 0.2 microns, followed by deconvolution and extended depth focus analysis to visualize 3D projections. Analysis was done using the Nikon GA3 processing tool, in which cells were defined in the DAPI channel. γH2AX, RAD51, pT21-RPA32, or pS33-RPA32 foci were quantified in an automated fashion for each cell using the Nikon GA3 BrightSpot function. Immunofluorescence staining of formalin-fixed paraffin-embedded tissue sections was performed as previously described including antigen retrieval, antibody conditions, and image acquisition, with no substantive modifications^39^.

### Xenograft studies

Xenograft studies were carried out under protocol no. 24-0237 approved by the Institutional Animal Care & Use Committee at the Washington University, School of Medicine. Six-week-old female NU/J mice were procured from Jackson Mice. Laboratory tumor implantation was performed once these mice were seven to eight weeks old. OVCAR8 cells were engineered to stably express a 2-in-1 SaCas9 plasmid expressing either sg*AAVS1* or sg*ALC1*. 3 × 10^6^ cells mixed 1:1 with Matrigel (Fisher Scientific, cat. no. 356237) were injected intraperitoneally into the mice. 14 days post-tumor implantation, two mice in each group were sacrificed to ensure the initiation of tumor nodules. 10 mice per group were randomized into vehicle- and drug-treated groups. Olaparib (selleckchem) was dissolved in captisol to a concentration of 50 mg ml^−1^. A 50 mg kg^−1^ dose of the drug was administered every second day for 35 days. At 35 days post-initiation of treatment, mice were sacrificed and analyzed for tumor weights and nodules.

### Statistics

Traditional statistical analyses were performed in GraphPad Prism 10 and SPSS version 27 software. OS was measured from time of PARPi initiation to date of death or last contact if no death was observed. Patients without recurrence or still alive were censored at their last contact date. Survival analyses were performed using Kaplan-Meier methods. Log rank test was used for group comparisons and Cox proportional hazards regression was used for estimating hazard ratios.

## Contributions

P.V. and M.M.M. conceived the idea and secured the funding. L.N.A and D.H.W. pursued most of the experiments, N.R. S.F.G, R.R. and D.K. assisted in qualitative immunofluorescence imaging, K.E.J. and V.M. helped with immunoblotting, and K.Z. helped with cloning under the supervision of P.V. E.L. and A.C. performed the xenograft experiments and helped with

IHC studies under the supervision of M.M.M. Dineo provided the TMA. E.L.C. analyzed the patient proteomics data. P.V. and M.M.M. wrote the manuscript with contributions from all other authors.

## Acknowledgements

We thank Dr. Alessandro Vindigni and the Genome Engineering and iPSC Center (GEiC) at the Washington University in St. Louis for the generation and validation of BRCA1 het FT282 cell lines. UWB SYr12 and SYr14 cell lines were obtained from Dr. Lee Zou at Duke University, School of Medicine. JHOS-4 and UWB-RR cell lines were obtained from Dr. John Krais and Dr. Neil Johnson at Fox Chase Cancer Center. This work was supported by the Inaugural Pedal the Cause Grant by Alvin J. Siteman Cancer Center through The Foundation for Barnes-Jewish Hospital, V-Scholar grant, Early-career investigator Award from Ovarian Cancer Research Fund Alliance, Career Catalyst Grant from Susan G. Komen and R37-CA286908 to P.V., who is also supported by Early-career Investigator Award from Ovarian Cancer Academy, Department of Defense, The Victoria’s Secret Global Fund for Women’s Cancers Career Development Award, in partnership with Pelotonia and the AACR and pilot project grant from Longer Life Foundation. L.N.A. is supported by the NIH Cancer Biology Pathway training T32 grant to Washington University, St. Louis and S.F.G. is supported by the NIH Cell and Molecular Biology training T32 grant to Washington University, St. Louis. MMM reports funding from the Damon Runyon Research Foundation, the Reproductive Scientist Development Program (RSDP) supported by the Gynecologic Oncology Group Foundation, the NCI Early-stage surgeon scientist program, the Victoria Secret Global Fund for Women’s Cancers Career Development Award in partnership with Pelotonia and AACR, the Pilot Translational and Clinical Studies function of the Washington University Institute of Clinical and Translational Sciences, and the Foundation for Barnes-Jewish Hospital, the Foundation for Women’s Cancer, and the American Cancer Society. M.M.M reports personal fees from Valinor Discovery and Bioscent Diagnostics.

