## Supplementary data and Tables for "Targeting ALC1 can safely expand the therapeutic utility of PARP inhibitors across high-grade serous ovarian cancers"

### Supplementary Materials

**Supp. Fig. 1**

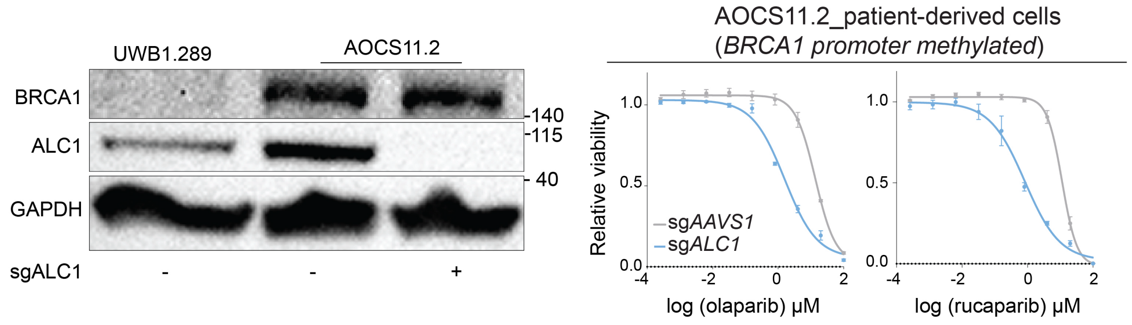


**Supplementary Figure 1. Loss of ALC1 enhances PARPi sensitivity in cells derived from ovarian tumor with *BRCA1* promoter methylation.**

Immunoblot (left) showing ALC1 depletion levels and PARPi sensitivity (right) of the patient-derived cells with BRCA1 promoter methylation quantified using the CellTiter-Glo assay; n = 3 biologically independent experiments. Data are presented as mean ± s.e.m.

**Supp. Figure 2**

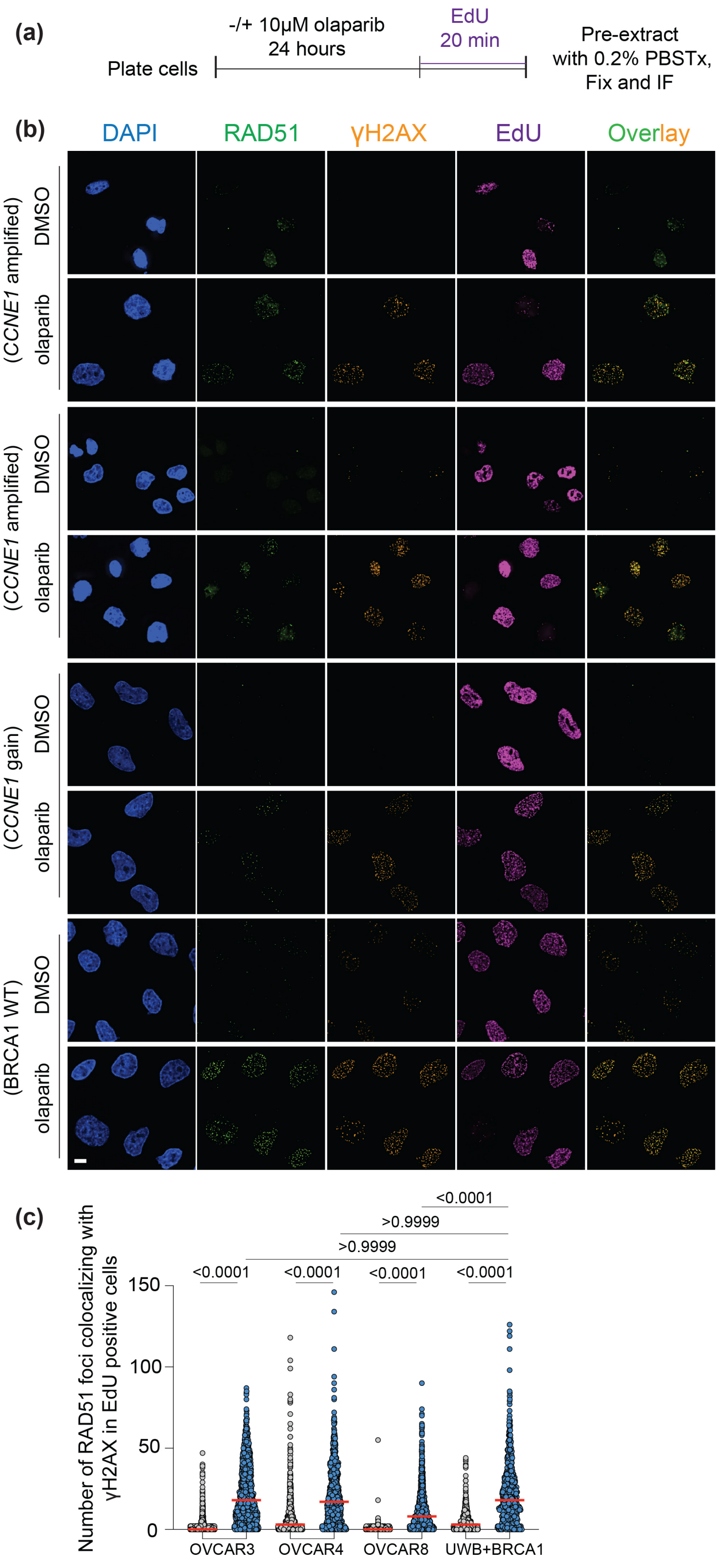

**Supplementary Figure 2. Cyclin E1-high ovarian cancer cells are proficient in loading RAD51 at DNA breaks post olaparib treatment.**

**(a)** Schematic of the experimental strategy to assess RAD51 loading post olaparib treatment.

**(b)** Representative images showing RAD51-gH2AX co-localized foci in indicated cell lines. Scale bar 10 microns.

**(c)** Quantification of RAD51-gH2AX colocalized foci in EdU positive cells of indicated cell line. Data are from n=3 biologically independent experiments; Median indicated. P-value: Kruskal-Wallis's test.

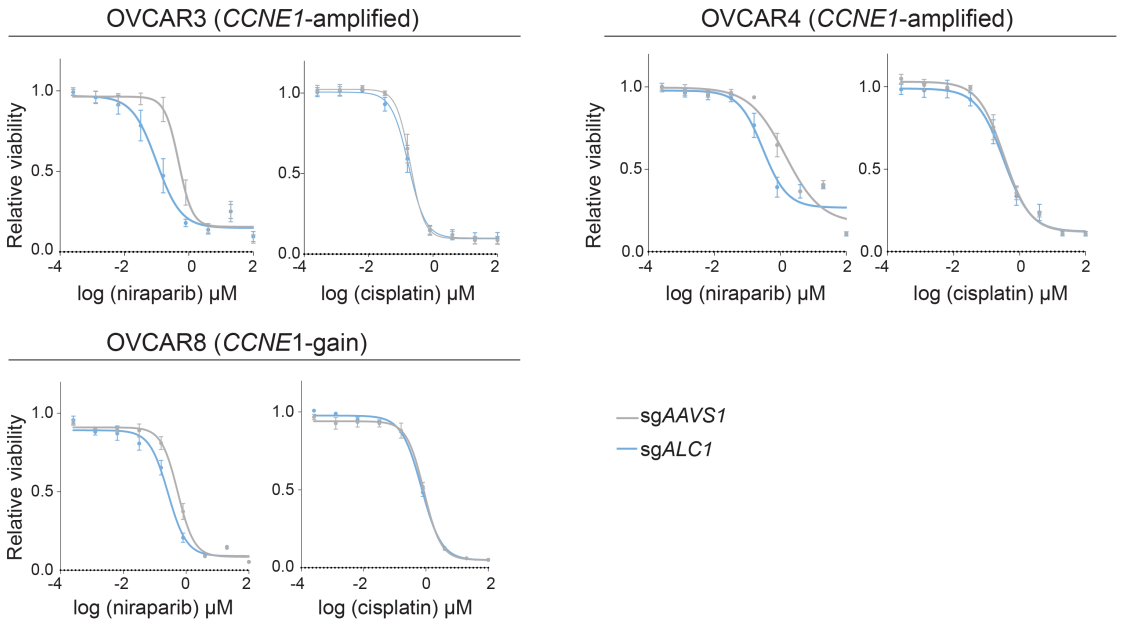


**Supplementary Figure 3. ALC1 loss results in minimal and no enhancement in sensitivity to niraparib and cisplatin respectively in cyclin E1-high serous cancer cell lines.**

Niraparib and cisplatin sensitivity of the indicated Cyclin E1-high ovarian cancer cells quantified using the Resazurin assay; n = 3 biologically independent experiments. Data are presented as mean ± s.e.m.

Supp. Figure 4

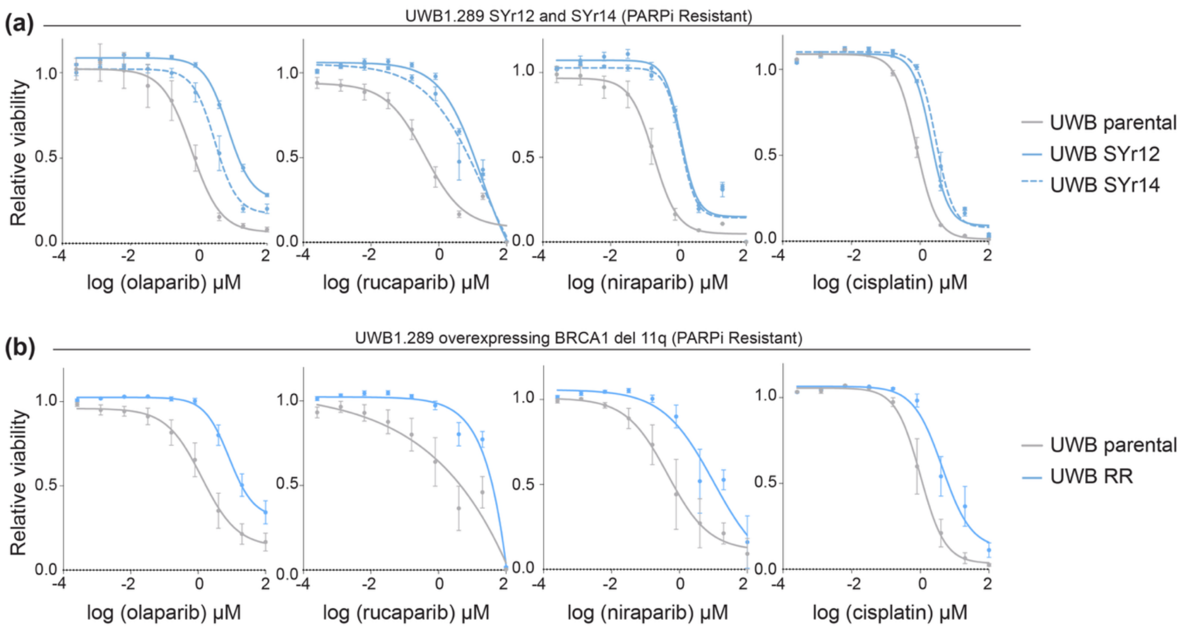


**Supplementary Figure 4. Characterization of PARPi resistant UWB1.289 ovarian cancer cells**

**(a)** PARPi and platinum sensitivities of the Syr12 and 14 UWB cell lines using the CellTiter-Glo assay; n = 3 biologically independent experiments. Data are presented as mean ± s.e.m.

**(b)** PARPi and platinum sensitivities of UWB cell line overexpressing del 11q using the CellTiter-Glo assay; n = 3 biologically independent experiments. Data are presented as mean ± s.e.m.

hTERT FT 282 {hTERT; p53 R175H; *BRCA1*+/c.222-223insA;(p.Glu75ArgfsX5)_(*BRCA1* het clone 1)

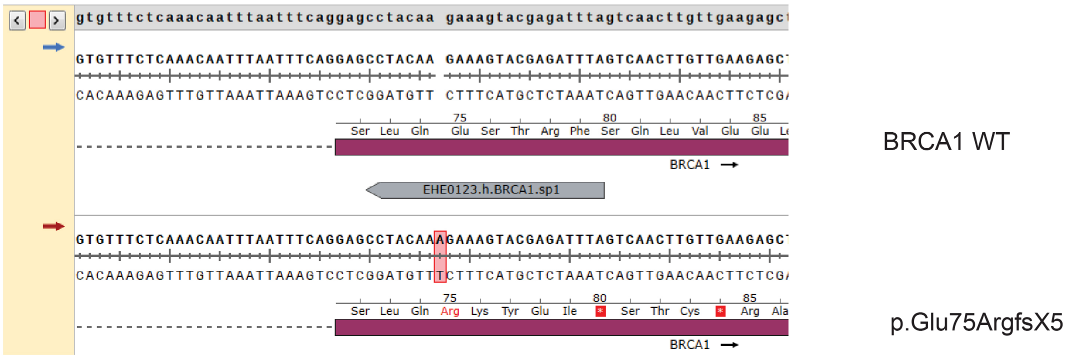


hTERT FT 282 {hTERT; p53 R175H; *BRCA1*+/c.223-226delAAAG;(p.Glu75ValfsX12)_(*BRCA1* het clone 2)

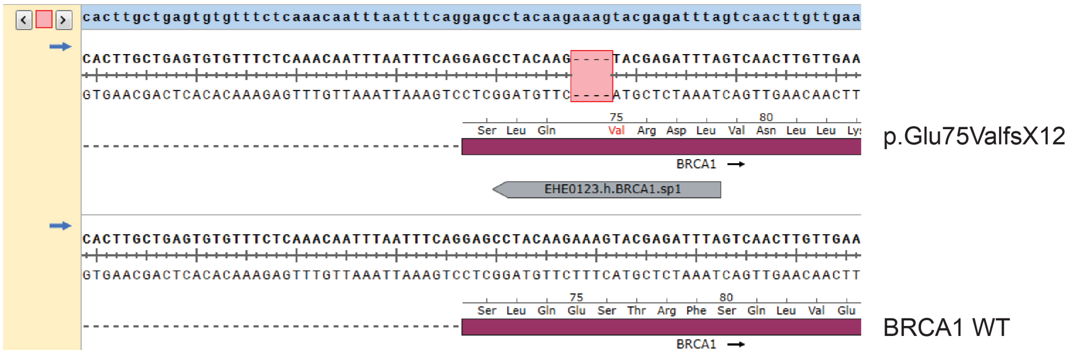


**Supplementary Figure 5. Generation of BRCA1+/- cell line.**

Sequencing data showing the *BRCA1* heterozygous mutations in the two clones derived from hTERT FT282 cell line.

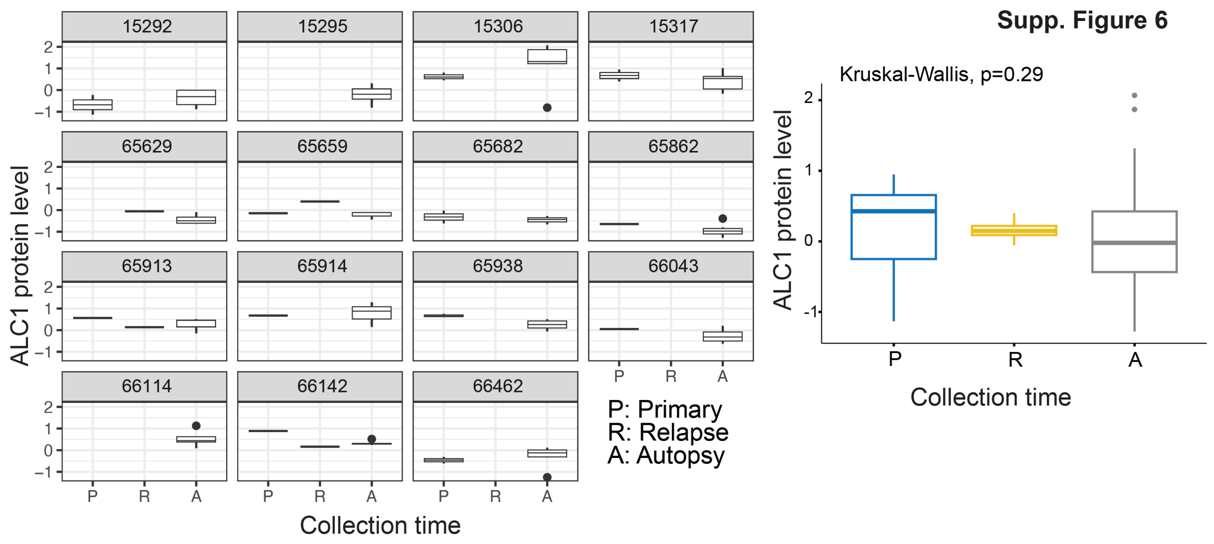


**Supplementary Figure 6. ALC1 protein level do not change over the course of PARPi treatment.**

Analysis of ALC1 protein levels in primary (P), relapse (R) and autopsy (A) samples separated by individual cases (left) and cumulative summary (right). The case number corresponds to the IDs from Burdett *et al*, 2023.

| Case | Primary | Relapse | Autopsy |
|---|---|---|---|
| 15295 | 0 | 0 | 7 |
| 65629 | 0 | 1 | 5 |
| 65862 | 1 | 0 | 8 |
| 65913 | 1 | 1 | 5 |
| 65914 | 1 | 0 | 3 |
| 65938 | 3 | 0 | 6 |
| 66114 | 0 | 0 | 7 |
| 66142 | 1 | 1 | 5 |
| 66462 | 2 | 0 | 7 |

**Supplementary Table 1.** Number of tumor samples collected from 9 individuals on PARPi. Case number refers to the IDs in Burdett *et al*[14]. Proteomics data from this paper was analyzed to assess correlation between ALC1/CHD1L protein levels and estimated time on PARP inhibitors.

**Primary Antibodies**

| Product | Catalog # | Lot # | Dilution used for western |
|---|---|---|---|
| Alpha-Actinin (D6F6) XP Rabbit monoclonal antibody | 6487S, Cell Signaling | 4 | 1:1000 |
| BRCA1 (D-9) mouse monoclonal IgG2a | sc-6954, Santa Cruz | G2821 | 1:250 |
| CHD1L(2170C3a) SAMPLE mouse monoclonal IgG1 | sc-81065, Santa Cruz | H1721 | 1:1000 |
| Anti-Cyclin E1 antibody [CCNE1/2460] mouse monoclonal IGg2b | Ab238081, Abcam | GR3437020-2 | 1:1000 |
| GAPDH (14C10) rabbit monoclonal | 2118S, Cell Signaling | 16 | 1:1000 |
| Alpa/Beta Tubulin polyclonal antibody | 2148S, Cell Signaling | 8 | 1:1000 |

**Secondary Antibodies**

| Product | Catalog # | Lot# | Dilution |
|---|---|---|---|
| ECL Anti-mouse IgG, Horseradish Peroxidase linked whole antibody (from sheep) | NA931, Cytiva | 17479274 | 1:1000 |

| | | | |
|---|---|---|---|
| ECL Anti-rabbit IgG, Horseradish Peroxidase linked whole antibody (from donkey) | NA934, Cytiva | 17348043 | 1:1000 |

**Supplementary Table 2:** Details of antibodies used for immunoblotting
